# The Effects of Intraspecies and Interspecies Competition on Genetic Device Construction and Performance

**DOI:** 10.1101/2025.06.17.660092

**Authors:** Samantha Thompson, A. Robert Williams, Veronica Dill, Deven Marshall, Emily Sawyer, Mason Alexander, Lilah Rahn-Lee, Joseph De-Chung Shih

## Abstract

Future genetic devices expressed in microbes need to be tested in complex environments containing multiple organisms to expand their usability. We sought to determine if genetic parts designed for monoculture are predictable when used in co-culture by testing constitutive Anderson promoters driving expression of chromoproteins from a plasmid. In *Escherichia coli* monoculture a high copy number origin of replication causes stochastic expression regardless of promoter strength, and high Anderson promoter strength leads to selection for inactivating mutations, resulting in inconsistent chromoprotein expression. Medium and low strength Anderson promoters function more predictably in *E. coli* monoculture but experience an increase in inactivating mutations when grown in co-culture over many generations with *Pseudomonas aeruginosa*. Expression from native promoters instead of Anderson promoters can lead to stable expression in a complex wastewater culture. Overall, we show intraspecies selection for inactivating mutations due to a competitive growth advantage for *E. coli* that do not express the genetic device compared to their peers that retain the functional device. We show additional interspecies selection against the functional device when *E. coli* is co-cultured with another organism. Together, these two selection pressures create a significant barrier to genetic device function in microbial communities that we overcome by utilize a native *E. coli* promoter. Future strategies for genetic device design in microorganisms that need to function in a complex microbial environment should focus on native promoters and/or strategies that give the microorganism carrying the device a selective or growth advantage.

**Importance:** First generation biotechnology focused on genetic devices designed for use in monoculture conditions. Next generation biotechnology devices will need to function in complex ecosystems with other organisms, so we sought to create conditions where the genetic device retained function when the organism carrying it is in co-culture with other organisms. We discovered that when the genetic device is a significant resource burden on the organism carrying the device mutations will be selected for due to intraspecies and interspecies selection pressures and the device will be rendered non-functional. Therefore, genetic device design for complex ecosystems in next generation biotechnology needs to balance functionality of the genetic device with the need to reduce resource burden on the organism carrying it.

## Background

Starting with the expression of human insulin in *E. coli* in the 1970’s [1], there has been an evolution in the design of genetic devices. The first generation of biotechnology utilized biological parts, such as promoters, terminators, and coding sequences, that were designed to maximize biotherapeutic or industrial yield in controlled monoculture environments such as bioreactors [2]. To expand beyond this model for biotechnology into more complex environments such as natural ecosystems and the human body, the next generation of genetic devices need to be designed for and tested in communities consisting of multiple microbial species.

One example of a field that could use such answers for next generation genetic devices is bioremediation. Microorganisms have been discovered in many different environments with natural bioremediation properties, such as the ones that were enriched during the 2010 Deepwater Horizon oil spill that were shown to degrade oil [3], microorganisms that degrade plastics found in the ocean [4], and others that sequester heavy metals in the soil [5]. These naturally occurring microorganisms found in the above studies have some bioremediation capability, but are slow in their natural rates of bioremediation, and so possess only limited natural bioremediation efficacy. To enable bioremediation at a scale and rate that would be environmentally useful, there are ongoing efforts to either genetically modify these organisms to increase their bioremediation rates or express combinations of enzymes important in bioremediation from naturally occurring but genetically intractable organisms in more genetically tractable organisms [6, 7]. However the organisms are made to enable scalable bioremediation, it is very likely that these organisms will have to exist in natural environments as a member of complex microbial communities to perform their designed functions.

We need to understand how the microbial community is affecting the genetic device and vice versa. Several questions will need to be addressed to achieve this goal. First, can a genetically modified organism enter an existing community and establish itself as a member? Genetically modified organism establishment has been demonstrated with using nutrient selection, such as in the case of Bacteriodes engineered to consume porphyrin fed as a prebiotic [8], but using such a specific nutrient limits potential utility. The rules that govern how genetically modified organisms establish themselves in a microbial community without such specific nutritional limits are being explored with in vitro and *in vivo* studies. [9–11] Second, does the introduction of the genetic device affect microbial community structure or function? Studies in soil microbe communities have shown that genetic device introduction can affect microbial community structure through changes in species abundance [12, 13], while functional changes in microbial communities caused by genetic device introduction have been modeled. [14] Finally, does the community impact genetic device function? We address this question in our study by putting a simple genetic device into microbial communities.

Therefore, we combined Anderson promoters, publicly available promoters with known performance capabilities [15], with commercially available chromoproteins from ATUM [16] in *Escherichia coli* and grew that *E. coli* with *Pseudomonas aeruginosa* in a simple co-culture community [11]. Anderson promoters and genes encoding chromoproteins are examples of genetic parts that could be particularly important for next generation genetic devices to operate in the environment, as constitutive promoters like Anderson promoters could be used to keep a genetic device on regardless of environmental conditions and chromoproteins could be useful as environmental reporters because they are detectable by the naked eye without need for a diagnostic device. These promoters and chromoprotein reporters have been individually optimized for expression in *E. coli* monoculture [15, 16], so assembling them as a genetic device in *E. coli* allowed use to test these parts in a microbial community. We determined that high strength constitutive Anderson promoters are poorly optimized devices in monoculture when expressed from high or medium copy number plasmids due to intraspecies selection for inactivating mutations and all Anderson promoters were poorly optimized in co-culture due to selection for inactivating mutations stemming from both intraspecies and interspecies competition. We suggest alternative methods for reliable expression of genetic devices undergoing both intraspecies and interspecies competition using native promoters and ribosome binding sites (RBS) that can function in co-culture.

## Methods

### Strains used in this study

*E. coli* strains DH5alpha and MG1655 were used where noted, and *P. aeruginosa* PAO1 was used for all co-culture assays. All bacteria were grown in LB Miller broth (Fischer Scientific, DF0446-70-5). *E. coli* DH5 alpha was purchased from New England Biolabs, *E. coli* MG1655 was purchased from Addgene.org, and *P. aeruginosa* was a gift from Dr. Stephen Lory. [17] For *E. coli* with pUC19 plasmids, 100 ug/mL ampicillin (Sigma, A9518) was used for selection, while for *E. coli* with pSB1C3 plasmids, 25 ug/mL chloramphenicol (Sigma, C0378) was used for selection. For overnight growth cells were grown in LB Miller broth shaking in 14 mL polystyrene snap-cap tubes (Corning, 352051) at 200 to 270 revolutions per minute and 37°C. For long-term assays cells were grown statically at 37°C in 5 mL of LB Miller broth in 14 mL polystyrene snap-cap tubes with 4-5 sterile glass beads [11]. Co-cultures of *E. coli* carrying pSB1C3 plasmids with *P. aeruginosa* were grown in LB media with chloramphenicol. To obtain roughly even starting co-culture cell concentrations, a 10:1 optical density at 600 nm (OD600) ratio of *E. coli* to *P. aeruginosa* was used for inoculation. Long term co-cultures and colony counts used methods from Ellis et al. [11]

### Device design and construction

IPTG-inducible ATUM chromoproteins on the pUC19 plasmid with ampicillin resistance were purchased from ATUM [16] and chemically transformed into *E. coli* DH5alpha cells. J23100 and J23108 Anderson promoters were ordered as primer dimers (Table 1) with AflII and XbaI restriction sites at the appropriate ends, PCR annealed using touchdown PCR from with denaturation at 95°C, annealing from 65°C to 55°C for 10 cycles followed by annealing at 55°C for another 25 cycles, and extension at 72°C , and inserted into the vector containing Tinsel Purple (tsP) and Cupid Pink (cuP) chromoproteins at AflII and XbaI restriction sites that cut out the original IPTG-inducible promoter using standard molecular biology techniques.

**Table 1:**
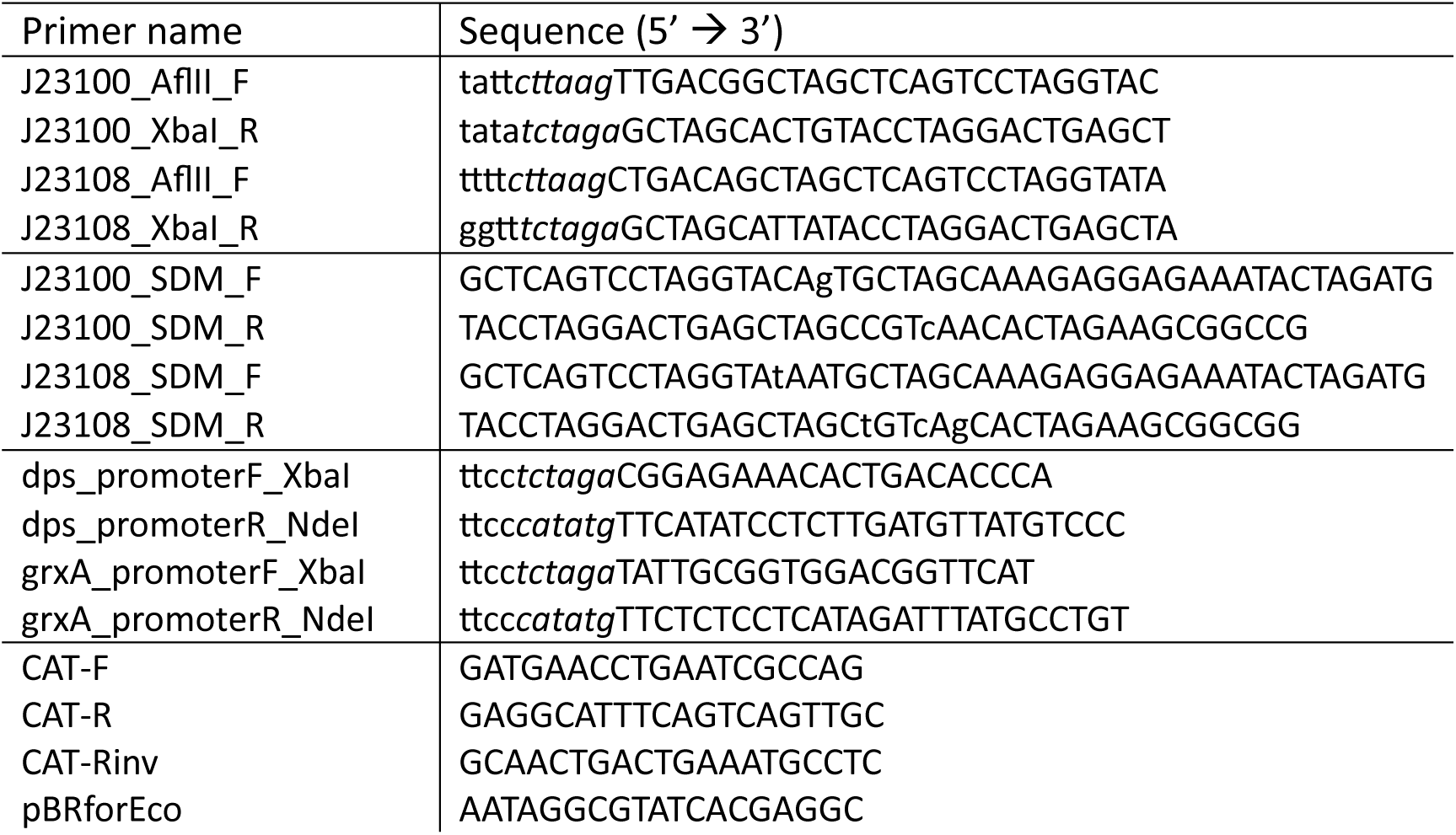
Primers used in this study for PCR amplification of operons and site directed mutagenesis (SDM). Italicized letters correspond to restriction sites. Capitalized letters correspond to sequences that will anneal to region being amplified.

tsPurple, a plasmid expressing tsP under the J23110 promoter, was a gift from Anthony Forster (Addgene plasmid # 117848) [18]. Site directed mutagenesis was performed using the Phusion High Fidelity PCR Kit (New England Biolabs) and SDM primers (Table 1) to mutate the J23110 promoter into the J23108 and J23100 promoters, DpnI (New England Biolabs) was used to digest the plasmid, and chemical transformation into *E. coli* DH5alpha cells was performed to produce plasmids JDS83 and JDS89, respectively. Successful mutagenesis was confirmed by Sanger sequencing using the pRBforEco primer (Table 1) [19]. tsPurple and JDS83 were chemically transformed into *E. coli* MG1655 for co-culture with *P. aeruginosa*.

To replace the Anderson promoters and strong RBS with promoters and RBS from the *dps* and *grx* [20] operons, dps_promoter and grxA_promoter primers (Table 1) were used to PCR amplify ∼400 bp of sequence upstream of those genes. The primers themselves contained XbaI and NdeI restriction sites that were used to insert the PCR products into the tsPurple plasmid at XbaI and NdeI restriction sites, cutting out the J23110 promoter and strong RBS using standard molecular biology techniques. The ligated product was chemically transformed into *E. coli* DH5alpha cells to produce plasmids JDS108 and JDS109 for the promoters and RBS from the *dps* and *grx* [20] operons, respectively. These plasmids were extracted using the ZymoPURE plasmid miniprep kit (Zymo Research) and chemical transformed into *E. coli* MG1655 for co-culture with *P. aeruginosa*.

### Diagnostic PCR and sequencing

Diagnostic PCR to detect presence of the chloramphenicol resistance (*cmr*) gene were done with primers CAT-F and CAT-R. Diagnostic PCR to detect the presence of the plasmid and to determine if deletions were present in the genetic device were done with primers pBRforEco [19] and CAT-R, and Sanger sequencing confirmation was conducted using primers pBRforEco and CAT-Rinv (Table 1). 1kb plus DNA ladder (N3200S) was purchased from New England Biolabs.

### Native cell lysis

Liquid cultures of each overnight culture were centrifuged at 5,000 g for 1 minute and supernatant was removed. Each cell pellet was resuspended in one volume of native cell lysis buffer consisting of 25 mM Tris-HCl pH 8, 50 mM sodium chloride, 2 mM ethylenediaminetetraacetic acid (EDTA), 1 mM dithiothreitol (DTT), 1 mM phenylmethylsulfonyl fluoride (PMSF), and 1 mg/mL of lysozyme from chicken egg white (Millipore), and incubated for 90 to 180 minutes at 37°C. The reaction was then centrifuged at 17,000 g for 5 minutes and the supernatant was removed and placed into cuvettes for spectrophotometry.

### Spectrophotometry and data analysis

Visible wavelength spectrophotometry from 400 nm to 800 nm was conducted on a Varian Cary 50 UV-Vis spectrophotometer (Agilent) with resolution of 1 nm using LB media as a blank for whole cell measurements or native cell lysis buffer without lysozyme as a blank for native cell lysis measurements. For whole cell measurements spectra were routinely above 1 OD in absorbance for the entire visible spectrum, so spectrometry data was aligned such that 800 nm = 1 OD for comparison purposes. For native cell lysis, spectrophotometry data was aligned such that each spectrum’s minimum value = 0 for comparison purposes. For chromoprotein peak calculations, a spectrum of *E. coli* MG1655 without chromoprotein expression was background subtracted. Statistical analysis of chromoprotein peak intensities was conducted using the emmeans package in R with the Tukey adjustment. [21]

### Mutant analysis

*E. coli* colonies containing plasmids with the tsP gene with no visible chromoprotein expression on LB agar plates with chloramphenicol selection were picked and grown overnight in LB media with chloramphenicol. An additional 20 ug/mL cefsoludin was added to the agar plates and liquid cultures if the *E. coli* were from co-cultures with *P. aeruginosa* to prevent *P. aeruginosa* contamination. Colonies were counted as not expressing tsP chromoprotein if there was no visible color after overnight incubation at 37°C and 2 additional days at 4°C. Plasmid DNA was isolated using the ZymoPure Plasmid Miniprep Kit (Zymo Research) and Sanger sequenced using either the pBRforEco sequencing primer [13] or CAT-Rinv (Table 1).

### Device testing in a wastewater microbial community

Wastewater was collected from the Liberty Wastewater Treatment Facility in Liberty, Missouri from the effluent prior to any treatment. LB media was inoculated with wastewater, then increasing concentrations of chloramphenicol were added and OD600 and percentage of colonies expressing tsP were measured to determine the appropriate concentration for the long-term assay. Every two days cell pellets were collected, colony PCR was conducted using pBRforEco and CAT-R primers to detect the presence of the plasmid, and on days 6 and 12 cell pellets underwent native cell lysis and spectrophotometry. Three biological replicate cell pellets each of wastewater grown overnight in LB media, grown 12 days in LB media, grown overnight in LB with 10 ug/mL chloramphenicol, and grown 12 days in LB with 10 ug/mL chloramphenicol, co-cultured overnight with *E. coli* MG1655 containing the JDS108 plasmid, and co-cultured for 12 days with *E. coli* MG1655 containing the JDS108 plasmid were collected and sent to Genewiz from Azenta Life Sciences for 16S rRNA metagenomic sequencing (16S EZ service). Amplicon sequence variants (ASVs) were inferred using dada2 [22], and taxa were assigned to ASVs using the Silva training set version 138.2 [23]. ASV abundances were graphed with phyloseq [24].

## Results

### ATUM chromoprotein expression is quantifiable

To determine if chromoprotein expression was robustly quantifiable, we first measured the visible absorbance spectra of eleven IPTG-induced chromoproteins expressed from the lac operon promoter on the pUC19 plasmid [25] obtained from ATUM [16] in *E. coli* and compared it with the visible absorbance spectrum of *E. coli* MG1655 without the plasmid. While all *E. coli* expressing chromoproteins formed pellets with qualitatively visible chromoprotein color as advertised by ATUM, their quantifiable absorbance peaks were highly variable. Of the purple chromoproteins, Tinsel Purple, Vixen Purple, and Prancer Purple had obvious absorbance peaks in the expected 580 – 590 nm range that had 0.27 – 0.54 greater absorbance than the *E. coli* MG1655 background (Figure 1A), while Cupid Pink had the largest absorbance peak of the non-purple chromoproteins in the 530 – 535 nm range that was 0.28 – 0.32 greater absorbance than the *E. coli* MG1655 background (Figure 1B). The large purple peaks found in Tinsel Purple, Vixen Purple, and Prancer Purple also had secondary peaks or shoulders at shorter wavelengths.

**Figure 1:**
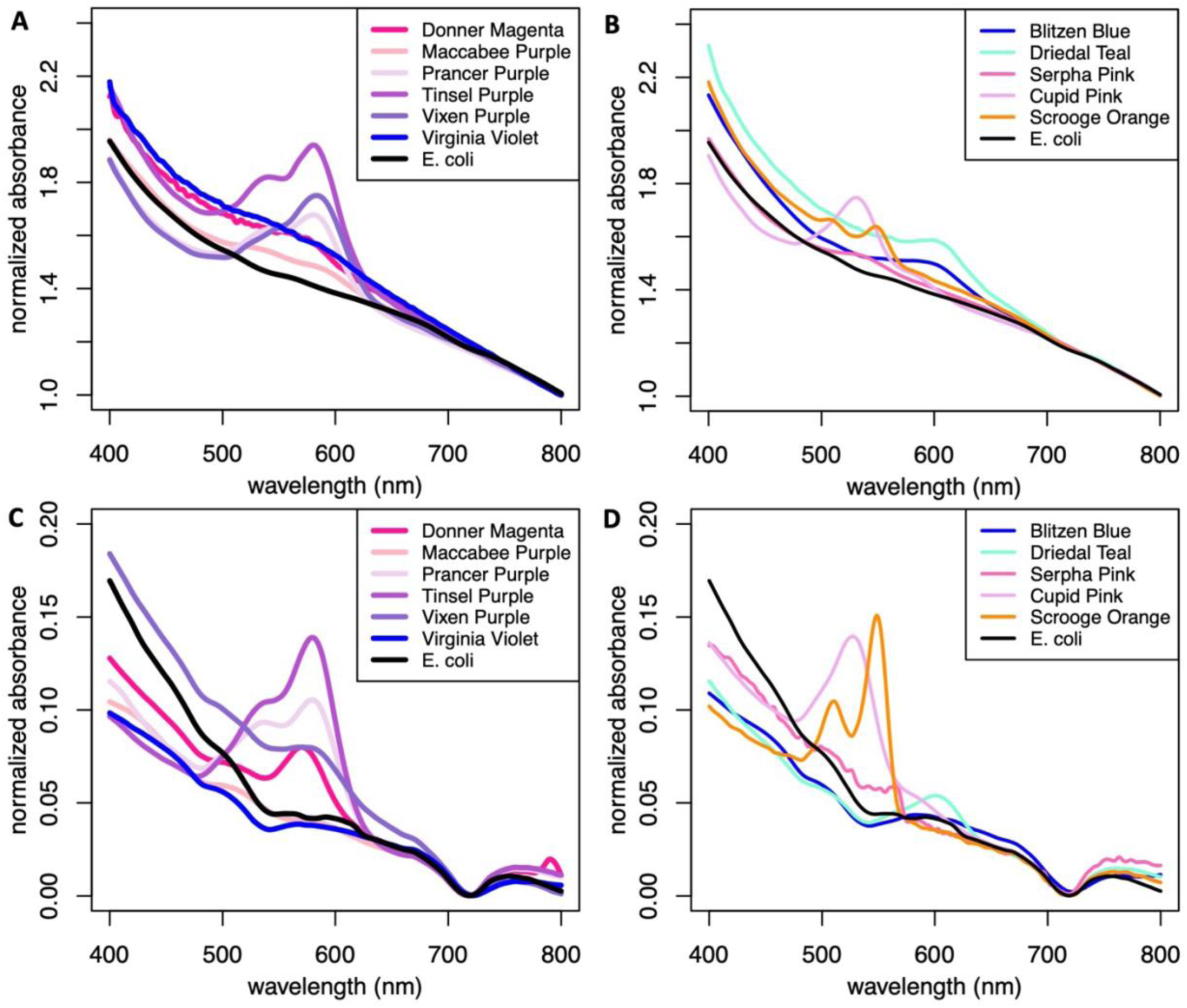
Chromoprotein absorbance spectroscopy. Absorbance spectroscopy of eleven IPTG-inducible chromoproteins grown in *E. coli* purchased from ATUM [16]. Native cell lysis of chromoproteins to preserve protein structure followed by pelleting and removal of cellular debris reduces background for measuring chromoprotein expression in monoculture. **A)** Purple chromoproteins in whole cell spectroscopy. **B)** Other chromoproteins in whole cell spectroscopy. **C)** Purple chromoproteins after native cell lysis. **D)** Other chromoproteins after native cell lysis.

Small peaks that had 0.07 – 0.16 greater absorbance than the *E. coli* MG1655 background were also noted for the rest of the purples and for Blitzen Blue, Dreidel Teal, and Serpha Pink, while Scrooge Orange had two small peaks at 505 – 515 nm and 545 – 555 nm. We conclude that chromoprotein expression is quantifiable in cells, as all the chromoproteins produced quantifiable primary peaks, but due to the variability of secondary peaks and shoulders we restrict our analysis to the primary peaks.

To reduce the high background we observed in whole-cell spectroscopy measurements, we performed native cell lysis to remove insoluble materials that scatter light, such as the lipid membrane and cell wall polymers, and measured the clear cell lysates after insoluble material was pelleted in the visible wavelength spectrum. Native cell lysis did reduce background, but it was a mixed success at keeping chromoproteins intact and functional as determined by absorbance peaks of the lysates. Of the purple chromoproteins, Tinsel Purple, Prancer Purple, and Donner Magenta benefited from native cell lysis, with peaks that had 0.05 – 0.1 greater absorbance than the *E. coli* MG1655 background, Vixen Purple’s absorbance spectrum had a reduced peak after native cell lysis than in whole-cell measurements, while the already small peaks for Maccabee Purple and Virginia Violet were eliminated (Figure 1C). In the non-purple chromoproteins Cupid Pink and Scrooge Orange had peaks with 0.08 – 0.11 greater absorbance than the *E. coli* MG1655 background, Dreidel Teal and Serpha Pink had peaks with only 0.01 –0.02 greater absorbance than the *E. coli* MG1655 background, and the peak for Blitzen Blue was eliminated (Figure 1D). Although native cell lysis did reduce the peak intensities of all the chromoproteins to varying degrees, it also reduced the background so significantly that it allowed for more precise measurements of the highest peak intensities, validating this method for use in chromoprotein peak measurements.

### Origin of replication and promoter strength affect successful performance of Anderson promoter driven chromoprotein genetic devices

Based on the highest peak intensities in visible wavelength absorbance spectra of native cell lysates, Tinsel Purple (tsP) and Cupid Pink (cuP) were chosen as the chromoproteins to be paired with Anderson promoters. We replaced the IPTG-inducible lac operon promoter on the pUC19 plasmid with constitutive Anderson promoters J23100 and J23108 (relative measured strengths = 1 and 0.51, respectively) [15], while retaining the strong T7 phage gene 10 RBS [26] (Figure 2A). We were successful in producing those constructs as confirmed by chromoprotein expression during cloning and subsequent Sanger sequencing but saw highly variable chromoprotein expression across clonal isolates in both liquid cultures grown overnight under chloramphenicol selection and on plates under chloramphenicol selection in all our constructs regardless of predicted Anderson promoter strength. This was true for constructs containing both tsP (Figure 2B) and cuP (Figure 2C) chromoproteins, demonstrating that it is difficult to have a reliable construct with predictable chromoprotein expression from the pUC19 plasmid.

**Figure 2:**
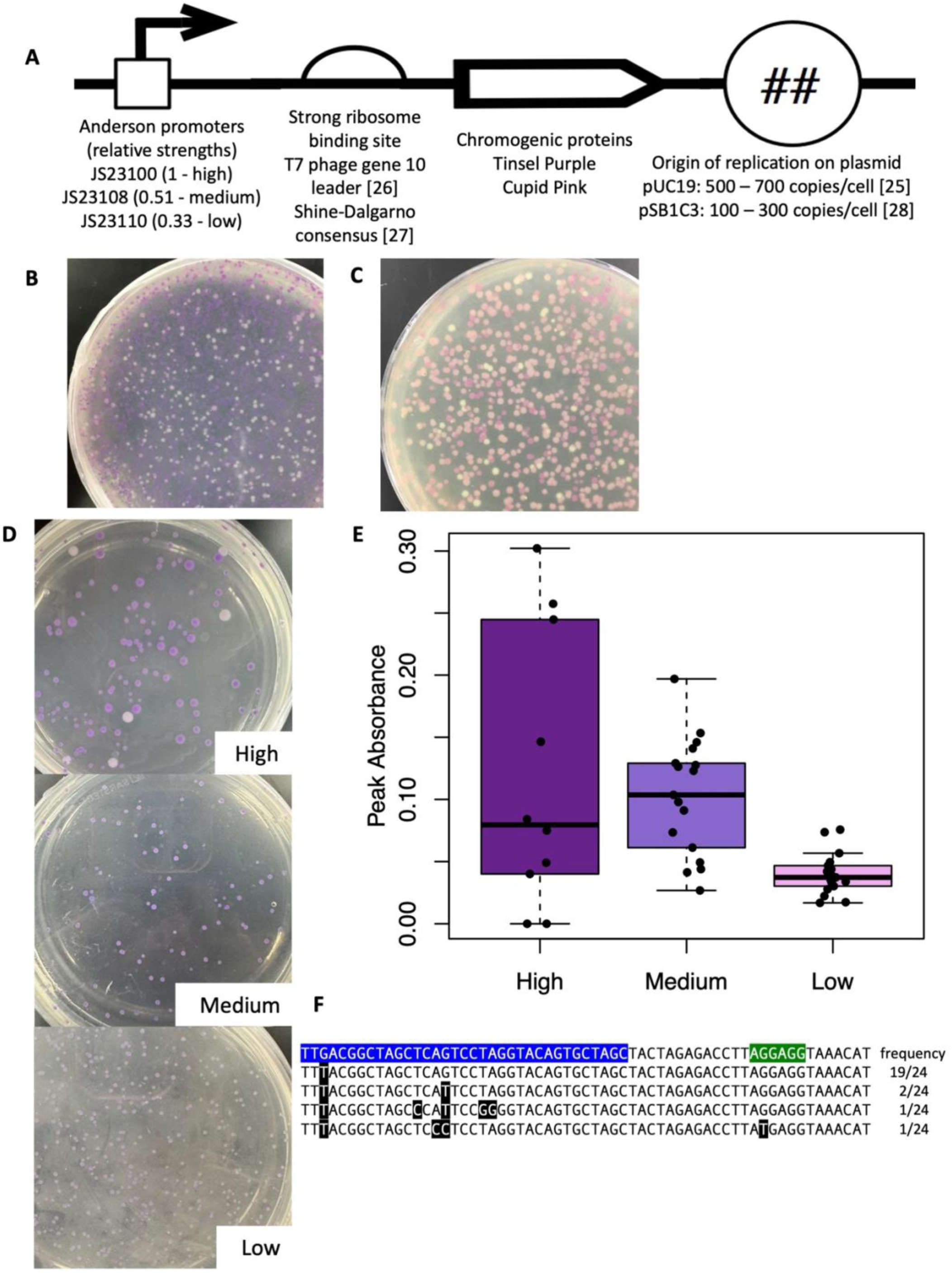
Chromoprotein expression is highly variable under high copy number origin of replication or high strength promoter. **A)** Device design. Anderson promoters of varying strengths were combined with genes encoding tsP or cuP chromoproteins on pUC19 [21] or pSB1C3 [24] plasmids. The strong RBS on each of the plasmids was unchanged throughout [22, 23]. **B)** Spread plate of high strength promoter driving tsP chromoprotein expression on the pUC19 plasmid. **C)** Spread plate of high strength promoter driving Cupid Pink chromoprotein expression from the pUC19 plasmid in *E. coli* DH5alpha cells grown over night in LB media under antibiotic selection. **D)** Spread plates of J23100 (high, top), J23108 (medium, center), and J23110 (low, bottom) promoters driving tsP chromoprotein expression from the pSB1C3 plasmid. **E)** Stable and predictable chromoprotein expression at lower promoter strengths from the pSB1C3 plasmid. Box and whisker plots of 580nm – 590nm peak absorbance values of tsP chromoprotein expression driven by Anderson promoters after overnight growth with antibiotic selection followed by cell lysis with background subtraction. Predicted relative promoter strengths are in parenthesis. N = 10 different colony peaks for high, 16 peaks for medium and low. **F)** Point mutations in the high strength Anderson promoter (blue) and Shine-Dalgarno consensus RBS (green) in mutants with low or no tsP expression. The top line is the original sequence, and other lines are sequences found in the 24 mutants isolated, with point mutations highlighted in black with white text.

We observed inconsistency in chromoprotein expression in supposedly clonal strains. We hypothesized that inactivating mutations were being selected for due to the very high copy number origin of replication on the pUC19 plasmid incurring a high resource cost [16]. This hypothesis was supported in a study by Liljerumn et al. [18], who produced stable tsP chromoprotein expression under the control of the J23110 Anderson promoter and a strong consensus prokaryotic RBS [27], but from the pSB1C3 plasmid, which has a relatively lower copy number origin of replication [28]. We obtained this plasmid, named tsPurple, and used site-directed mutagenesis to mutate the J23110 promoter (relative measured strength = 0.33, low strength) into the J23100 and J23108 promoters (relative measured strengths = 1 and 0.51, high and medium strength, respectively) [15], then spread plated each of these constructs after overnight liquid culture, with both plates and liquid culture nder chloramphenicol selection (Figure 2D). From these spread plates we picked and grow up 10 – 16 single colonies in liquid culture under chloramphenicol selection, performed native cell lysis, measured the absorbance spectra, and quantified the maximum peak absorbance values after subtracting the *E. coli* MG1655 absorbance spectrum background (Figure 2E). The high strength promoter had the most variability of tsP chromoprotein expression amongst the individual colonies, with some clonal isolates not producing any chromoprotein expression after native cell lysis. The high strength promoter was not significantly different from the medium strength promoter. It did have significantly greater peaks compared to the low strength promoter (P = 0.0058), but we determined that it was too variable to be used in future assays. Conversely, both the medium strength and low strength promoters combined with tsP expressed on the pSB1C3 plasmid in *E. coli* produced clonal isolates with predictable tsP chromoprotein peaks in the 580nm – 590nm range as quantified by absorbance spectra of native cell lysates. While the medium strength promoter produced more variable peak intensities than the low strength promoter, it had reliably greater peaks than the low strength promoter (P = 0.0139), leading us to conclude that these promoters driving tsP expression were reliable enough to be used in future assays.

We suspect that the high-strength promoter was producing selection against tsP chromoprotein production due to high metabolic resource usage, much like the high copy number origin on pUC19 did, and therefore mutations occurring in the promoter or coding sequence that disabled or reduced chromoprotein production would provide a fitness advantage that was selected for over time. We grew five independent inoculations of *E. coli* with the high strength Anderson promoter driving tsP chromoprotein expression in monoculture over twelve overnight passages in static conditions, plated dilutions of the final culture, screened for *E. coli* colonies with no visible tsP chromoprotein despite chloramphenicol selection, and Sanger sequenced the promoter, ribosome binding site, and tsP coding sequence (CDS) of 24 of these isolated colonies from across the five independent inoculations (Figure 2F).

All the isolates had at least one point mutation in the promoter, with a few that had many different point mutations in the promoter and RBS, supporting our hypothesis that mutations that reduced or disabled chromoprotein production conferred a growth advantage that was selected for over time. The most common point mutation that all the isolates had was the G to T transversion in the third nucleotide of the promoter. This transversion is found in 8 out of 18 Anderson promoters that have reduce measured promoter strength compared to the strongest Anderson promoter, [15] suggesting that this position influences promoter strength. Some isolates had mutations in the middle region that is constant among most of the Anderson promoters, which would likely lead to loss of promoter function due to the strong sequence conservation found that region of Anderson promoters. [15]

### Growth of Anderson promoter driven genetic devices in co-culture with *P. aeruginosa* eliminates chromoprotein expression in *E. coli* by producing selection against functional devices

Since tsP expression from the pSB1C3 plasmid driven by the medium and low promoters had predictable expression in *E. coli* MG1655 monoculture, we decided to co-culture *E. coli* MG1655 containing each device with *P. aeruginosa* PAO1, as it is well-established that *E. coli* MG1655 and *P. aeruginosa* PAO1 can exist in stable co-culture in static conditions [6, 29–31]. We grew this co-culture under chloramphenicol selection to keep the plasmid in *E. coli,* as *P. aeruginosa* PAO1 is naturally chloramphenicol resistant [32]. We found that hosting neither genetic device changed the ability of *E. coli* to participate in co-culture with *P. aeruginosa,* (Figures 3A and 3B) as colony forming units (CFU) per mL (CFU/mL) were comparable with prior co-culture results seen by Ellis *et al.* [11] We could not detect any tsP expression in liquid culture, either by eye or by native cell lysis and spectrophotometry at any point in our co-culture beyond the original day of inoculation (Figure 3C). Monocultures of *E. coli* with the genetic devices being grown statically and passaged for the same amount of time as the co-cultures reliably expressed tsP (Figure 3C). One possible explanation for not detecting tsP in co-culture is that the presence of cell lysate material from *P. aeruginosa* could interfere with the absorbance of tsP, while another explanation is that tsP is not being expressed in co-culture. To determine which of these possibilities is true, we mixed statically grown monoculture populations of *E. coli* and *P. aeruginosa* at similar ratios as seen in long-term co-culture and immediately conducted native cell lysis on them (Figure 3D). While the peaks in the lysate of mixed monocultures were lower and the background higher than in *E. coli* monocultures, the peaks were visible in these controls, suggesting that interference by *P. aeruginosa* cell lysate debris alone cannot account for the complete lack of tsP peaks in long-term co-cultures. This result suggests that the co-culture prevents some or all tsP expression either by reducing transcription and/or translation of tsP, or by selecting for mutations that inactivate the functional device.

**Figure 3:**
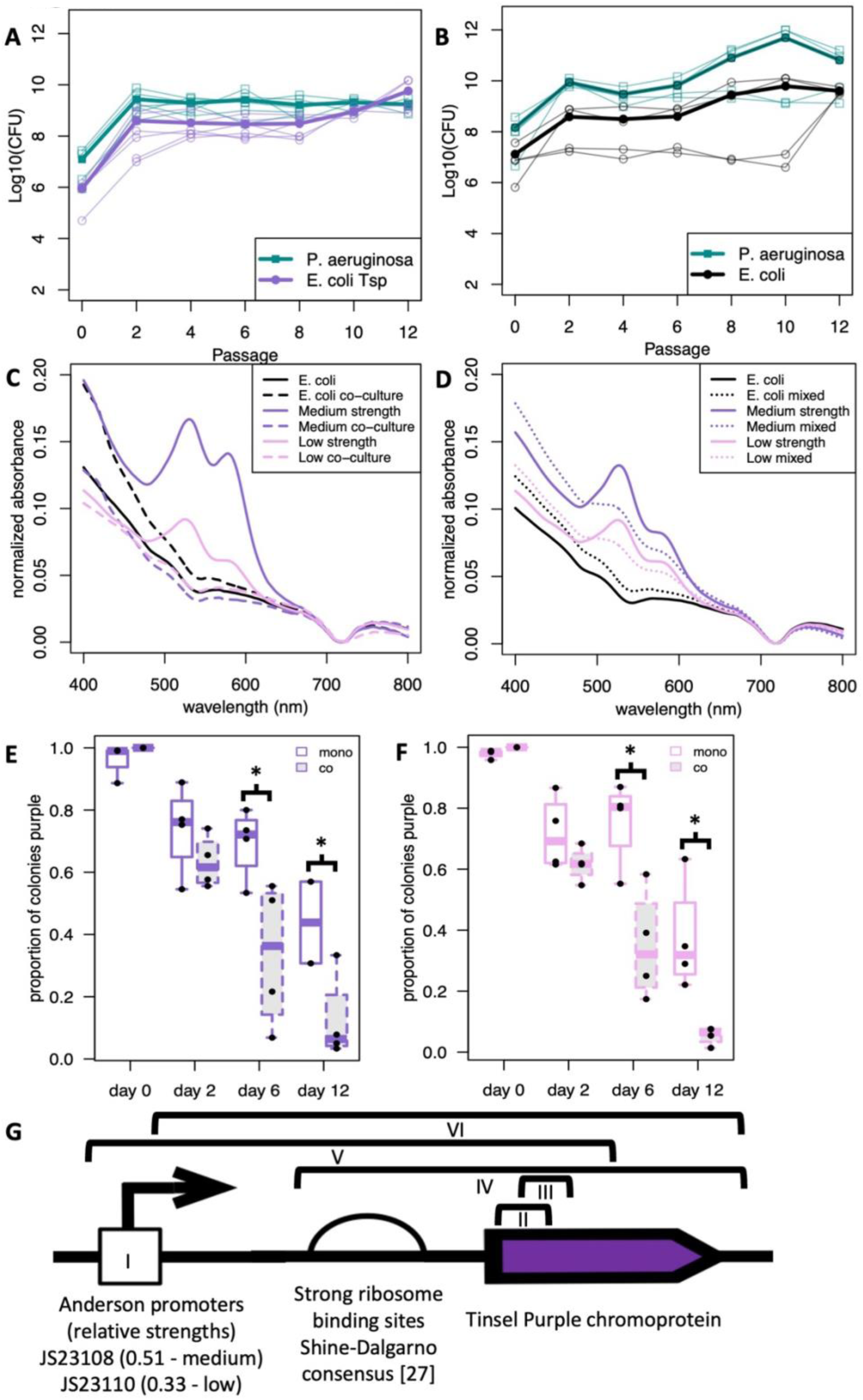
Genetic devices in E. coli co-cultured with P. aeruginosa does not alter community stability, but results in no tsP chromoprotein expression. **A)** Log 10 of colony forming units per mL for *E. coli* MG1655 expressing medium or low strength Anderson promoters driving tsP expression (purple) co-cultured with *P. aeruginosa* PAO1 (green). N = 6. Thin lines show individual experiments, thick lines show the average. Each passage is a liquid culture grown overnight, dilution plated, and counted after overnight incubation at 37°C. **B)** Colony counts for *E. coli* MG1655 (black) co-cultured with *P. aeruginosa* PAO1 (green). N = 4. Thin lines show individual experiments, thick lines show the average. Each passage is a liquid culture grown overnight, dilution plated, and counted after 24 hours growth at 37°C. **C)** E. coli expressing tsP grown in co-culture with *P. aeruginosa* PAO1 has no detectable tsP expression. The medium strength promoter driving tsP expression in monoculture (solid purple) and in co-culture with *P. aeruginosa* PAO1 (dashed purple) and the low strength promoter driving tsP expression in monoculture (solid pink) and in co-culture with *P. aeruginosa* PAO1 (dashed pink) were passaged every day for 10-12 days before cells were lysed and supernatant absorbance was measured. *E. coli* MG1655 monoculture (solid black) and *E. coli* MG1655 co-cultured with *P. aeruginosa* PAO1 (dashed black) were included as negative controls. **D)** P. aeruginosa cell debris presence in lysate cannot account for complete loss of tsP detection in co-cultures. *E. coli* MG1655 expressing tsP under medium (solid purple) and low (solid pink) strength Anderson promoters were grown in monoculture and native cell lysis and spectroscopy was performed on either the monoculture or a 1:1 cell number mixture with *P. aeruginosa* PAO1 (dotted purple and dotted pink). *E. coli* MG1655 monoculture (black) and mixed with *P. aeruginosa* PAO1 (dotted black) were included as negative controls. Proportion of *E. coli* colonies carrying functional devices expressing tsP from **E)** medium and **F)** low strength Anderson promoter in monoculture after days of co-culture with *P. aeruginosa* is lower than in monoculture over time. Each day is one passage. N = 4, error bars = standard deviation. * P < 0.05 comparing monocultures to co-cultures at days 6 and 12, Student’s T-test. **G)** Qualitative summary of mutations found in medium and low strength Anderson promoters and Shine-Dalgarno consensus RBS in 32 mutants with no tsP expression. Sections indicated by the black brackets are deletions. The mutations are: (I) point mutations in the promoter similar to those found in the high strength promoter, (II) 79 bp frameshift deletion in tsP (2 – 80) resulting in an early STOP after 46 amino acids, (III) 78 bp in-frame deletion in tsP resulting in a 26 amino acid deletion, (IV) deletion from the end of the promoter to beyond the end of tsP, (V) deletion of the promoter, RBS, and first 400bp of tsP, (VI) deletion after first 14 bp of the promoter to beyond the end of tsP.

To look for mutations that inactivate the functional device, we tested whether the device was functional by plating liquid co-cultures on plates with cefsulodin selection, which excludes *P. aeruginosa*, and chloramphenicol selection, which retains the device,. Since a colony on a plate is a physiological condition in which functional devices would produce tsP, any white colonies on these plates would be due to mutants that formed and were selected for during co-culture. Over time we observed a decrease in *E. coli* colonies that showed expression of tsP after co-culture with *P. aeruginosa* compared to *E. coli* grown in monoculture despite antibiotic selection on plates for the plasmid (Figures 3E and F). As a control we removed chloramphenicol selection on plates, resulting in more rapid decline of tsP expression in the *E. coli* colonies than on plates with chloramphenicol selection for both monocultures and co-cultures (Supplemental Table 1). This control argues against plasmid loss as the cause of most of the lost tsP expression in *E. coli* co-cultured with *P. aeruginosa*. Much like the high strength promoter is producing selection against tsP expression in *E. coli* grown in monoculture (Figure 2E), we conclude that *E. coli* in co-culture with *P. aeruginosa* is producing increased selection against tsP expression compared to *E. coli* monoculture in the medium and low strength promoters, and therefore selecting for inactivating mutations occurring in the promoter or coding sequence over time.

Sanger sequencing of the promoter, RBS, and tsP CDS of 22 *E. coli* colonies with devices with medium or low strength Anderson promoters with no tsP expression after co-culture with *P. aeruginosa* for 12 days and in 10 *E. coli* colonies grown in monoculture for 12 days all showed either point and/or deletion mutations. (Figure 3G, Supplemental Table 2). The point mutations were in the conserved middle of the Anderson promoters, similar to prior mutations seen in the high strength Anderson promoters (Figure 2F), and the deletions removed some or all the promoter region and some or all the tsP coding sequence. Two deletions were frameshift deletions within the tsP coding sequence. One deletion is predicted to produce a truncated 46 amino acid peptide, while the other deletion is an in-frame deletion, but an Alphafold 3 prediction of the mutant [33] produces an in-frame deletion removes the majority of the first strand and all of the second strand of the beta barrel, causing incomplete beta barrel formation and suggesting a loss of function due to loss of beta barrel integrity.

### A genetic device utilizing a native promoter results in detectable expression in co-culture

While Anderson promoters may only be suitable for *E. coli* monoculture, it will be desirable for genetic devices that interact with microbial communities to have high expression either only in communities or in both communities and in monoculture. In long-term co-culture of *E. coli* MG1655 with *P. aeruginosa* PAO1 [11], we used RNA sequencing to discover genes in *E. coli* that were more highly expressed in co-culture with *P. aeruginosa* than in monoculture. We selected *dps* and *grxA* as genes with high base expression in monoculture and higher expression in co-culture than monoculture, with *dps* having roughly an order of magnitude more expression than *grxA* in both scenarios (Figure 4A). We asked if genetic devices could be optimized for co-culture by utilizing promoters and RBS from these genes. We replaced the Anderson promoters and the consensus strong RBS with promoters and RBS from the *dps* and *grx* operons in *E. coli* MG1655 [20] while keeping the tsP chromoprotein and origin of replication on the pSB1C3 plasmid intact (Figure 4B), creating *P_dps_-tsP* and *P_grxA_-tsP* devices, respectively. We found that neither genetic device in *E. coli* significantly changed the proportion of *E. coli* in co-culture with *P. aeruginosa*, there was no statistical difference in CFU/mL with *E. coli* MG1655 co-cultured with *P. aeruginosa* and prior co-culture results [11] (Supplemental Figure 1). Monoculture of *E. coli* with both devices expressed tsP (Figure 4C), with *P_dps_-tsP* producing much greater peaks than *P_grxA_-tsP*, in line with their baseline *E. coli* expression (Figure 4A). In long-term co-culture with *P. aeruginosa*, *E. coli* with the *P_dps_-tsP* device produced small but detectable amounts of tsP, while *E. coli* with the *P_grxA_-tsP* device failed to produce any detectable tsP in co-culture with *P. aeruginosa* using the cell lysis assay (Figure 4C). Monocultures of statically grown populations of *E. coli* with the *P_dps_-tsP* device mixed with P*. aeruginosa* at similar ratios as seen in long-term co-culture with native cell lysis produced reduced but still visible peaks (Figure 4D), similar to what was seen in long-term co-culture (Figure 4C), suggesting that the *P_dps_-tsP* device functioned as well in co-culture with *P. aeruginosa* as in monocultures. Both native promoter devices we used experienced selection against functional devices across our 12 day assay (Figures 4E and F). *E. coli* with *P_dps_-tsP* had an increased proportion of colonies expressing tsP when co-cultured with *P. aeruginosa* from day 6 to day 12 (Figure 4E), suggesting either sampling bias or drift. Monocultures of *E. coli* with either device started with high percentages of colonies expressing tsP, but by day 12 those percentages were similar to the percentages of colonies in co-culture with *P. aeruginosa,* suggesting similar expression levels and selection for inactivating mutations in co-culture and monoculture.

**Figure 4:**
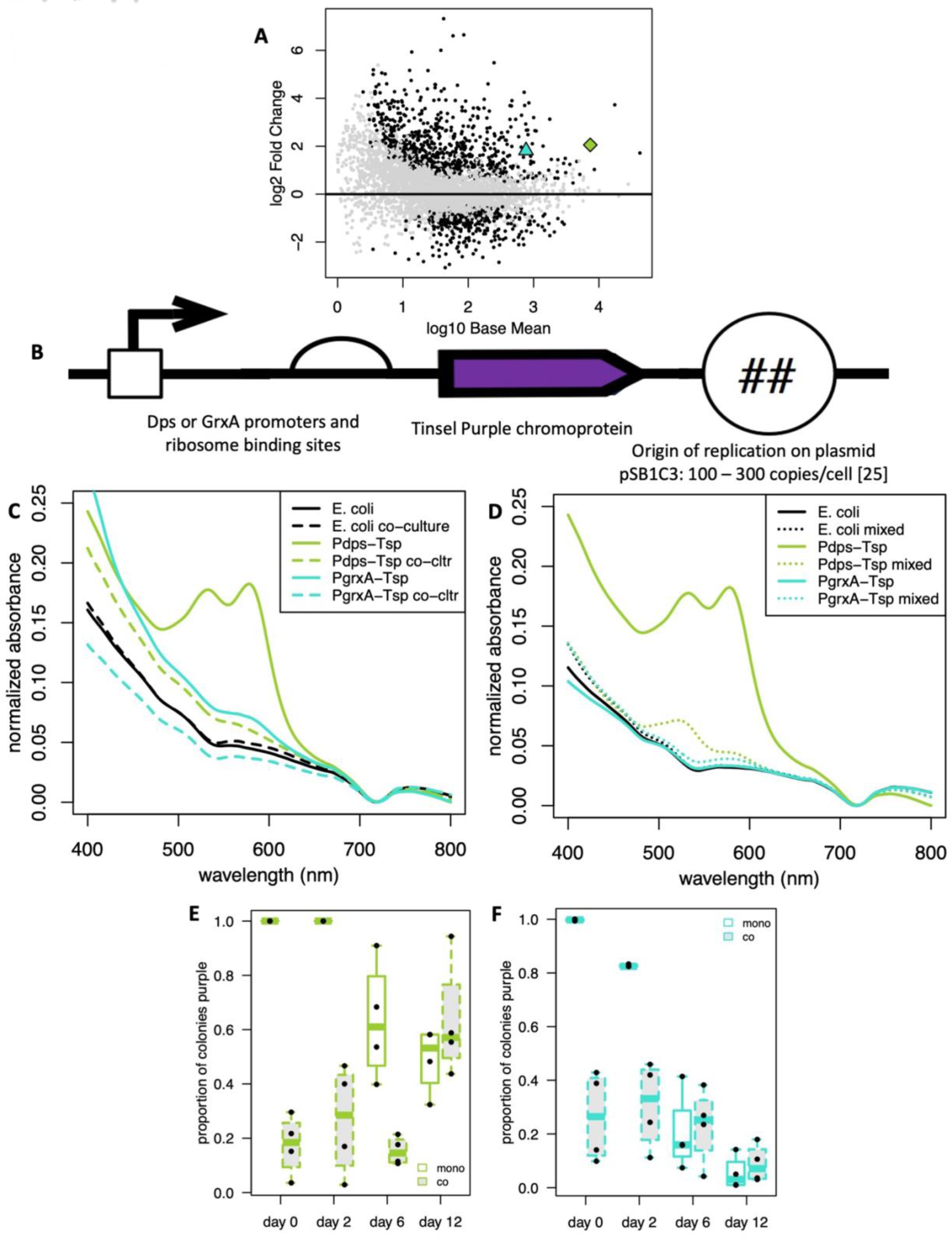
Native promoters optimized for increased expression in E. coli *during co-culture with* P. aeruginosa produce detectable expression in co-culture. **A)** RNA sequencing comparing log10 base mean gene expression of *E. coli* MG1655 in monoculture and log2 fold change in expression after 8 – 12 days of co-culture with *P. aeruginosa* PAO1. Black dots are individual *E. coli* genes that are significantly upregulated (above 0) or downregulated (below 0) in co-culture compared to monoculture. Green diamond = *dps*, Blue triangle = *grxA*. **B)** Device design. *dps* or *grxA* promoters and RBS were combined with the gene expressing the tsP chromoprotein on the pSB1C3 plasmid. **C)** tsP expression of *E. coli* expressing native devices grown overnight in monoculture (*P_dps_-tsP* = solid green and *P_grxA_-tsP* = solid blue) or co-cultured for 12 days with *P. aeruginosa* PAO1 (*P_dps_-tsP* = dashed green and *P_grxA_-tsP* = dashed blue) detected through native cell lysis and absorbance spectrophotometry. *E. coli* MG1655 monoculture (solid black) and *E. coli* MG1655 co-cultured with *P. aeruginosa* PAO1 (dashed black) were included as negative controls. **D)** E. coli with *P_dps_-tsP* has detectable tsP expression when grown in monoculture, then mixed with *P. aeruginosa*. *E. coli* MG1655 with native devices were grown in monoculture and native cell lysis and spectroscopy was performed on either the monoculture (*P_dps_-tsP* = solid green and *P_grxA_-tsP* = solid blue) or a 1:1 cell number mixture with *P. aeruginosa* PAO (*P_dps_-tsP* = dashed green and *P_grxA_-tsP* = dashed blue). *E. coli* MG1655 monoculture (solid black) and *E. coli* MG1655 co-cultured with *P. aeruginosa* PAO1 (dotted black) were included as negative controls. Proportion of *E. coli* colonies with **E)** P_dps_-tsP or **F)** _PgrxA-tsP_ in co-culture with *P. aeruginosa* expressing tsP. Each day is one passage. N = 4.

At the end of our long-term assay with the native promoter devices, we isolated plasmid DNA from 61 *E. coli* colonies with no visible tsP expression, 24 from the *P_dps_-tsP* device and 21 for the *P_grxA_-tsP* device grown in co-culture with *P. aeruginosa* for 12 days and 8 *E. coli* colonies grown in monoculture for 12 days for each native promoter device, and conducted diagnostic PCR reactions to detect the presence of various parts of the plasmid. PCR with primers internal to the *cmr* gene on the plasmid produced mostly positive hits, with 16/17 diagnostic reactions, 9 for *P_dps_-tsP* and e8 for *P_grxA_-tsP*, producing the expected 511 bp band (Figure 5A), and the one not producing the expected 511 bp band producing a slightly bigger band, indicating retention of the *cmr* gene in most colonies. Sanger sequencing of colonies from native devices that had no visible tsP expression when co-cultured with *P. aeruginosa* using the pBRforEco primer revealed some of the colonies had mutations and some did not, with 6/9 *P_dps_-tsP* and 1/14 *P_grxA_- tsP* colonies producing chromosomal integration events excluding the tsP CDS (Figure 5B, mutation II, Supplemental Table 3). Almost half of the Sanger sequencing reactions using pBRforEco produced no result, suggesting a lack of primer binding. Diagnostic PCR with pBRforEco and CAT-R of devices grown in co-culture with *P. aeruginosa* for 12 days revealed that 22/45 colonies, 15 from *P_dps_-tsP* and 7 from *P_grxA_-tsP*, did not produce any band at all (Figure 5C). Since the *cmr* gene was confirmed to be present (Figure 5A), we suspected plasmids from these colonies were missing the pBRforEco primer binding site, so we sequenced with CAT-Rinv, an inverted sequence of CAT-R primer found in the *cmr* gene which is predicted to be 227 bp upstream from pBRforEco. Sanger sequencing with CAT-Rinv resulted in the discovery of additional chromosomal integration events and deletions that resulted in loss of the pBRforEco primer binding site (Figure 5B, mutations I, IV, & V, Supplemental Table 3). Overall, 21/24 isolated *E. coli* colonies with *P_dps_-tsP* co-cultured with *P. aeruginosa* with no tsP expression had deletion mutations or random chromosomal integration events and 8/21 isolated *E. coli* colonies with *P_grxA_-tsP* co-cultured with *P. aeruginosa* had deletion mutations or random chromosomal integration events. Meanwhile, 3/8 isolated *E. coli* colonies with *P_dps_-tsP* grown in monoculture produced the same mutation (Figure 5B, mutation III, Supplemental Table 3), a deletion of most of the promoter and RBS and 78 bp of the tsP CDS with a small 15 bp insertion, while none of the *E. coli* colonies with *P_grxA_-tsP* grown in monoculture displayed a mutation.

**Figure 5:**
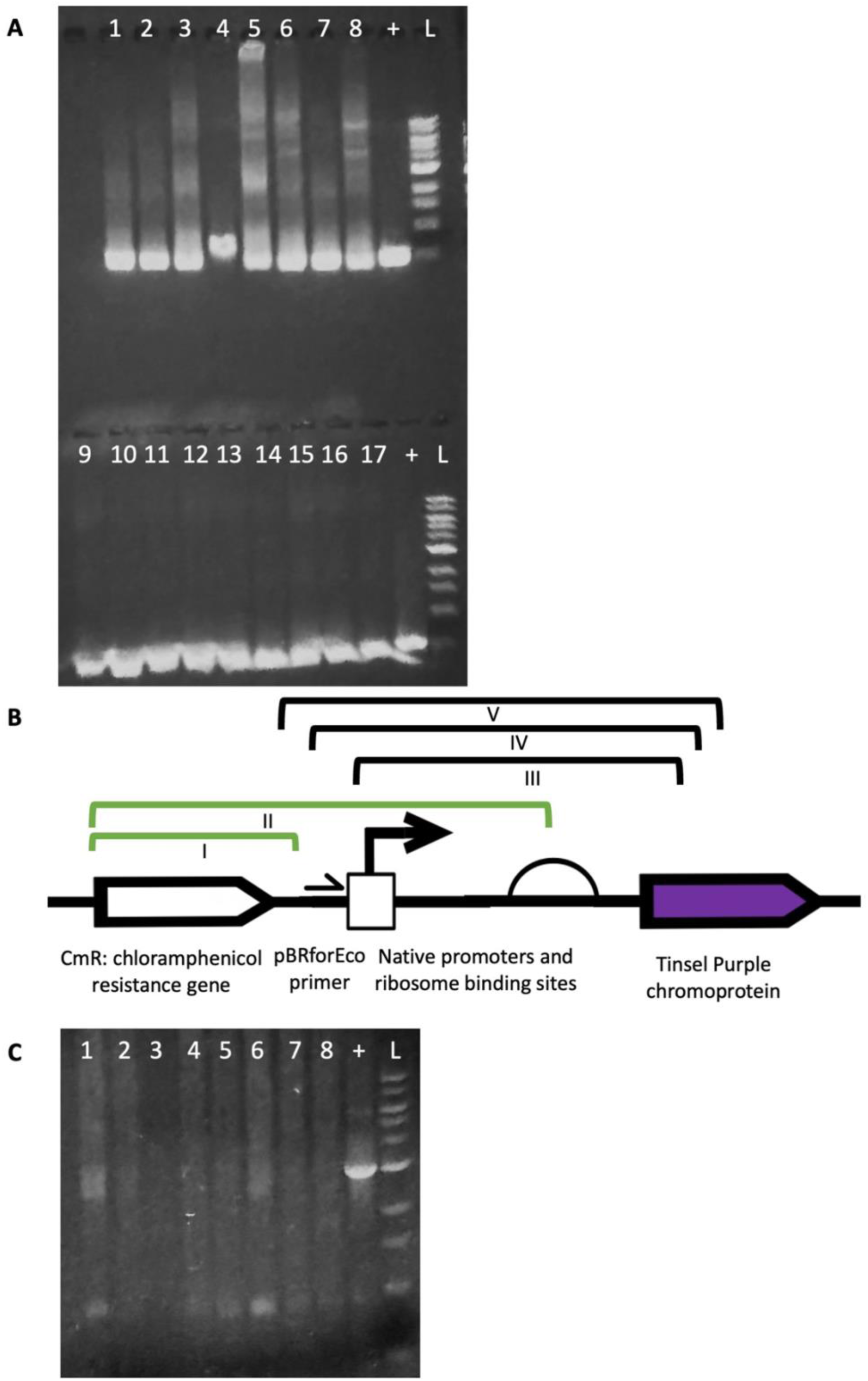
Intraspecies vs interspecies competition correlate with different types of mutations. **A)** A diagnostic PCR conducted to detect the *cmr* gene. Individual colonies of *E. coli* with *P_grxA_-tsP* device (1 – 8) and *Pdps-tsP* (9 – 17), + = tsPurple plasmid positive control, L = 1kb DNA ladder. **B)** Qualitative summary of mutations found in native promoter and RBS devices in 61 co-culture and monoculture mutants with no tsP expression. Sections indicated by the black brackets are deletions. Sections indicated by the green brackets are regions of the plasmid that have been randomly integrated into the *E. coli* chromosome. The mutations are: (I) Chromosomal integration of the *cmr* gene, (II) Chromosomal integration of the *cmr* gene and most of the promoter and RBS region (III) Deletion from 2bp into promoter ending after first 78 bp of tsP, with a 15 bp insertion (TGGTGGCAAGTTATG), (IV) Deletion starting at the pBRforEco primer ending after the first 329 bp of tsP, (V) Deletion starting 40 bp downstream of *cmr* ending after the first 394 bp of tsP. **C)** Representative diagnostic PCR to determine if the pBRforEco binding site was present. Individual colonies of *E. coli* with PgrxA-tsP (1 – 4) and Pdps-tsP (5 – 8), + = tsPurple plasmid positive control, L = 1kb DNA ladder

Since all the colonies we isolated had no visible tsP expression, this suggests that there are other mutations responsible for the phenotype, either elsewhere on the plasmid or in *E. coli* MG1655 genome, that we could not detect.

### A native promoter and RBS device is functional in a natural microbial community

Since *E. coli* with *P_dps_-tsP* functioned in co-culture with *P. aeruginosa,* we wanted to assess how it would perform in a natural community over time. To do this, we obtained untreated wastewater effluent from a local wastewater treatment facility and tested the device in the presence of diverse and abundant wastewater bacteria to determine if tsP would still be expressed and if it would impact the wastewater microbial community.

Our *P_dps_-tsP* device is on a plasmid with the *cmr* gene, so it is important that selection for the device is maintained with chloramphenicol. Therefore, we had to determine what concentration of chloramphenicol would have minimal effect on the natural community members of the wastewater while still maintaining selection for the device, which we experimentally determined to be 10 μg/mL (Figure 6A). We performed a modified long-term assay under 10 μg/mL instead of 25 ug/mL chloramphenicol selection for 12 days. Triplicate cell pellets of the wastewater co-cultured with *E. coli* with *P_dps_-tsP* were purple to the naked eye for all 12 days, with visible decrease in tsP expression by day 10. (Figure 6B) Plasmid extraction and diagnostic PCR using pBRforEco and CAT-R primers targeting the plasmid confirmed the plasmid was present (Supplemental figure 2), and spectrophotometry of the cell lysate on days 6 and 12 showed obvious tsP peaks (Figure 6D), leading us to conclude that our *P_dps_-tsP* device remained present and functional in the natural wastewater community under 10μg/mL chloramphenicol selection.

**Figure 6:**
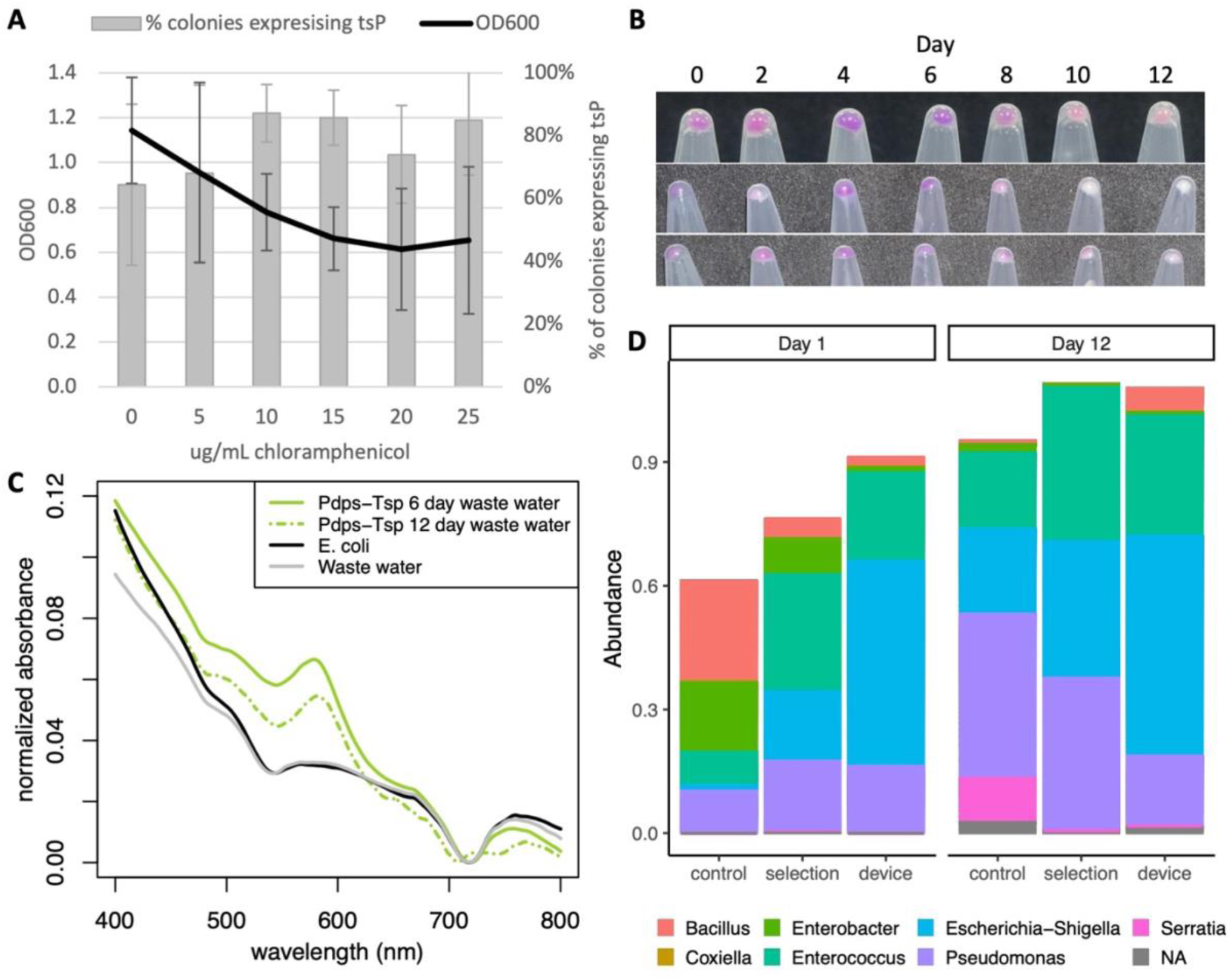
Expression of a native promoter device in a wastewater community. **A)** Selecting an optimal concentration of chloramphenicol to maintain plasmid selection and wastewater community. Wastewater microbes were grown overnight at in LB with various concentrations of chloramphenicol and OD600 measured after overnight growth. N = 6. *E. coli* MG1655 with *P_dps_-tsP* was grown overnight in LB with various concentrations of chloramphenicol, plated, and colonies were assayed for tsP expression. N = 3. **B)** Visible tsP expression in wastewater throughout the 12-day assay. Cell pellets were spun down every two days. Each row is a biological replicate. Cell pellets from left to right are day 0, 2, 4, 6, 8, 10, and 12. **C)** tsP expression of *E. coli* with *P_dps_-tsP* cultured for 6 days (solid green) and 12 days (dashed green) with wastewater community detected by native cell lysis and absorbance spectrophotometry. *E. coli* MG1655 monoculture (black) and wastewater community (gray) were included as negative controls. **D)** Normalized abundance of the genera of the most common 400 taxa identified by 16S rRNA sequencing from wastewater inoculum grown in LB (control), LB with 10 ug/mL chloramphenicol (selection), or LB with 10 ug/mL chloramphenicol and co-inoculated with E. coli carrying the *P_dps_-tsP* device (device), sampled after one or twelve days of serial passage.

We performed 16S rRNA metagenomic sequencing to determine what members were in the wastewater community and if our *P_dps_-tsP* device with chloramphenicol selection altered the community (Figure 6D). Chloramphenicol selection did enrich for some of the most common genera originally found in the wastewater after 1 day of growth compared to growth without chloramphenicol, but it merely accelerated enrichment that long-term growth in rich LB media promoted, as there was little difference in the abundance of the top three genera after 12 days for growth in LB with or without chloramphenicol selection or between these and 1 day of chloramphenicol selection. Adding *E. coli* with *P_dps_-tsP* increased the proportion of *Escherichia-Shigella* in both day 1 and day 12, as would be expected if *E. coli* with *P_dps_-tsP* were maintained as a significant portion of the microbial community, but did not significantly alter the frequency of the other top two genera in the microbial community grown with 10 μg/mL chloramphenicol in either day 1 or day 12, leading us to conclude that our functional device had minimal impact on wastewater community composition.

## Discussion

Our goal is to determine if robust genetic parts that are designed for predictable use in monoculture are as robust as claimed when put into complex microbial communities. This is particularly relevant considering the need for future genetic devices to function in complex microbial environments. Based on our findings, we conclude that current genetic devices optimized for monocultures are not robust when asked to perform the same functions in complex microbial communities. Conversely, at no point in our experiments did it seem that our genetic devices had any significant effect on community composition within a complex microbial community, either in *E. coli* co-cultured with *P. aeruginosa* (Figure 3A) or in the wastewater microbial community (Figure 6D). We cannot rule out changes in genetic expression and community function in organisms when a functional genetic device is present that could be detectable using transcriptional profiling, but complex microbial communities seem to be structurally robust in a way that genetic devices themselves are not.

While we acknowledge that the primary objective of ATUM was to create chromoproteins with qualitative visibility, it was surprising to us that some of them were difficult to quantify using whole cell absorbance spectroscopy in the visible wavelengths as the chromoprotein peaks were highly variable in size. This quantifiable variability was further amplified by our native cell lysis procedure, which did succeed in dramatically reducing background but at the cost of reducing chromoprotein peak intensities overall. While the largest peaks were more precise following native cell lysis, many of the small peaks were eliminated. Due to the proprietary nature of these chromoproteins, we do not know their protein structure and thus cannot speculate on specific reasons why one chromoprotein may have kept or lost its peak(s), but we can conclude that our attempt to preserve protein structure using our native cell lysis procedure needs refinement. There is likely some protein denaturation, and some modifications to this procedure in the future to reduce protein denaturation while preserving cell lysis could be considered, such as adjusting salt, pH, or buffer concentrations to preserve protein folding or increasing the amount of phenylmethylsulfonyl fluoride (PMSF) protease inhibitor.

We originally created devices with two different chromoproteins and two Anderson promoters of varying strengths on a plasmid with a high-copy number origin of replication. The chromoprotein expression of these devices as seen on plates was determined to be highly variable, with no predictable pattern. This variability led us to switch to a plasmid with a lower copy number origin of replication and focus on one chromoprotein, which produced predictable expression of tsP chromoprotein at lower Anderson promoter strengths. While the RBS were different on both plasmids, they were both previously known to be strong RBS [26, 27], so we chose not to alter those when constructing our devices to reduce the complexity of our study.

In addition to promoter strength and origin of replication, variation in RBS is another way to vary genetic device expression and could be the focus of future studies. We show that lowering the copy number by switching from pUC19 to pSB1C3 does provide some stability and predictability in genetic device function. One way to eliminate the influence of copy number on genetic device function would be to integrate genetic devices into the microbe’s genome and avoid plasmids entirely, thus providing stability with a low copy number that would result in less costly genetic device function. However, this would likely come with the trade-off of lower protein expression and therefore negatively impact genetic device functionality. As long as initial genetic device construction is conducted on plasmids, plasmid copy number will need to considered for device validation, and there could be situations when where it may be suitable to have the genetic device only temporarily in an organism, such as when the microbe naturally exists in the environment and the genetic device is intended for temporary function, like in immediate bioremediation.

Regardless of whether genetic devices are integrated into the genome, high promoter strength can interfere with reliable genetic device function. In our study, once the origin of replication strength was reduced, the Anderson promoter with highest promoter strength still demonstrated high variability in chromoprotein expression. Previous work has shown that high strength Anderson promoters lose expression upon entering a competitive microbial environment [34] and negatively affect *E. coli* growth [35], while in a prior study *E. coli* with the low strength Anderson promoter driving tsP expression from the pSB1C3 plasmid had 75% - 80% of the growth rate *E. coli* compared with a promoterless control plasmid [18]. Together, these results lead us to hypothesize that expressing genetic devices reduce growth rates in *E. coli,* but genetic devices on a high copy number plasmid or with the strongest Anderson promoter reduce growth rates to the point of inordinately selecting for mutations that inactivate tsP chromoprotein expression. Some of the tsP expression variability from the pUC19 plasmid could also be from selection for similar inactivating mutations in promoters, so it is likely that copy number and promoter strength additively contribute to the cost of a genetic device.

*E. coli* expressing our constitutive Anderson promoter genetic devices in co-culture with *P. aeruginosa* failed to express any measurable tsP chromoprotein, despite the equivalent *E. coli* monoculture producing robust chromoprotein expression and mixtures of monocultures also producing visible, albeit weaker, chromoprotein peaks. *E. coli* with *P_dps_-tsP* had high tsP expression in monoculture and low tsP expression in long-term co-culture with *P. aeruginosa,* while *E. coli* with *P_grxA_-tsP* had low tsP expression in monoculture and no tsP expression in long- term co-culture with *P.* aeruginosa (Figure 4C). We propose two explanations for these results.

First, there is a dampening effect from *P. aeruginosa* cell lysate, which we found in *E. coli* monoculture controls of both Anderson promoters expressing tsP (Figure 3D) and native devices (Figure 4D) mixed with *P. aeruginosa*. It is unknown how much of the lack of tsP chromoprotein expression in co-culture with *P. aeruginosa* is due to selection for mutants and how much is due to changes in gene expression of functional devices due to co-culture context, though detectable tsP expression in both day 6 and day 12 *E. coli* monocultures for Anderson and native promoter devices despite the presence of significant inactivating mutations suggests some of the lack of tsP expression in co-culture is due to changes in gene expression.

We observed an increase in inactivating mutations in co-cultures of *E. coli* with medium and low strength Anderson promoters expressing tsP with *P. aeruginosa* compared to monoculture (Figures 3E and F), suggesting that co-culture plays a role in selecting for inactivating mutations as it imposes additional interspecies selection pressures. *E. coli* is in a very different physiological and metabolic state in co-culture with *P. aeruginosa* compared to monoculture. It undergoes a starvation response and switches away from aerobic respiration [11], factors that could make gene expression more costly and therefore select for inactivating mutations. This suggests that while genetic devices always incur a cost, there is a cost threshold at which selection results in loss of functional devices from the population. In our work here, co-culture had a lower cost threshold, perhaps due to the added interspecies competition, than monoculture did.

It is interest to note that the type of mutations we saw differs by type of genetic device utilized. *E. coli* with medium and low strength Anderson promoters experience both point and deletion mutations regardless of whether they are in monoculture or co-culture (Figure 3G, Supplemental Table 2). This is different from the high strength promoter, where we saw only point mutations (Figure 2F). We saw that the high strength Anderson promoter imposed a large resource burden on *E. coli* causing loss of tsP expression in monoculture (Figures 2D and 2E) and wonder if that burden could explain the selection for point mutations. Surprisingly, we observed no point mutations in the inactivated native promoters. Point mutations are more common than other types of mutations, comprising 95% of mutations found in human solid tumors [36] and almost two-thirds of mutations in the SARS-CoV-2 Spike glycoprotein [37], and amongst other types of mutations deletions are more common than insertions [38]. Most insertions and deletions tend to be 50 bp or less [38], making the deletions and random integrations we see in our inactivated native promoters unusual. We speculate that these longer deletions and integrations are more effective in inactivating a genetic device but occur less frequently. Therefore, there are instances, such as utilizing the high strength Anderson promoter, where the selection pressure against the functional genetic device is so strong that the need to inactivate the device selects for more common point mutations as the mechanism of inactivation. Conversely, if the selection pressure is not as strong, then point mutations may not produce a strong enough phenotypic difference to warrant selection, so over time the mutations that are selected for are the more effective longer deletions and integrations that remove large portions of the genetic device.

We observed loss of function mutations arising in both Anderson devices and native promoter devices over time (Figures 3E and F, Figures 4F and G). However, the *P_dps_-tsP* device retained functionality in the native lysis assay (Figures 4C and 6C) and in the wastewater microbial community (Figure 6B). This suggests that *P. aeruginosa* lysate masking and loss of function mutations cannot completely explain why the Anderson devices do not function during co-culture. Ellis et al. [11] has shown that *E. coli* undergoes the starvation response when in co-culture with *P. aeruginosa.* Even though Anderson promoters are conceived of as constitutively expressed, this expression relies on the “housekeeping” σ^70^ sigma factor, which is modulated during the starvation response [39, 40]. Dps plays a role in protecting DNA during oxidative stress and starvation and is upregulated in during the starvation response [41]. The *dps* promoter is bound by σ^70^ , but also contains has a consensus σ^S^ binding site (GCTATACTTAA) in the -10 position [42, 43]. σ^S^ is the stress sigma factor, so it would likely be activated during co-culture with *P. aeruginosa* when *E. coli* is starving [11]. Therefore, *dps* and other genes that are activated by Sigma S [44] are good candidates for reliable promoters to drive expression of genetic devices in the context of co-culture with *P. aeruginosa* due to activation of the starvation response. Thus, to create a functional device it is important to consider the RNA transcript data and the regulatory environment of the gene, even when using nominally “constitutive” promoters.

Putting a genetic device in a microbe that provides no survival or growth benefit to that microbe while incurring a resource utilization cost introduces a selective pressure to reduce the expression of that genetic device because metabolic and gene expression resources that could be used for growth are diverted to genetic device function. This is a well-known phenomenon [45–47] further supported by reduced growth rates found in *E. coli* expressing chromoproteins [18] and by the increase in mutations selected for in co-cultures found during our current study.

Selective pressure can be a result of intraspecies competition within one species, as microbes with the functional genetic device are at a competitive disadvantage with same species microbes with a mutated nonfunctional or reduced function genetic device. Furthermore, when putting a microbe with a genetic device in an environment with different species the additional selective pressure of interspecies competition is introduced. This additional interspecies selective pressure has profound consequences, as even devices that seem to function reliably with low selection bias in monoculture may be stretched beyond their functional breaking point in a more complex microbial community. Our results support the need to test next generation genetic devices in model microbial communities to make sure they can function as designed before any large-scale environmental release.

## Conclusions

Based on our study, we reject the hypothesis that genetic devices and parts can have predictable functionality regardless of what other genetic parts they are coupled with or environmental context they are placed in. Future design and construction of robust genetic devices or parts for predictable functionality need to account for their environmental conditions and genetic context, not just for the organism they are to be expressed in. For complex microbial communities, we suggest not using constitutive promoters and using promoters that are optimized for expression in those communities. Constructing genetic devices that will give the microbe with that device a selective survival or growth advantage in a particular environment could also enable robust genetic device function.

## List of abbreviations

CDS: coding sequence
CFU: colony forming units
*cmr*: chloramphenicol resistance
DTT: dithiothreitol
EDTA: ethylenediaminetetraacetic acid
PMSF: phenylmethylsulfonyl fluoride
RBS: ribosome binding site
tsP: Tinsel Purple
cuP: Cupid Pink

## Declarations

### Ethics approval and consent to participate

Not applicable

### Consent for publication

Not applicable

### Availability of data and materials

All data generated and analyzed during this study are included in this published article. The datasets used and/or analyzed during the current study are available from the corresponding author on reasonable request.

### Competing financial interests

The authors declare that they have no competing interests. *Funding:* This study was funded by the U.S. National Science Foundation award 2031102. *Author’s contributions*: ST conducted the analysis of mutations for both co-cultures and monocultures, preparation of samples for Sanger sequencing, the long-term growth experiments for the native devices driving tsP in *E. coli* during co-culture with *P. aeruginosa*, co-culture of *E. coli* MG1655 containing the Pdps-tsP device with wastewater microbiome and contributed to writing the manuscript. ST and AW constructed the native promoters optimized for expression in *E. coli* during co-culture with *P. aeruginosa*. VD and AW conducted the experiments with the medium and low strength Anderson promoters driving tsP expression in *E. coli* in co-culture with *P. aeruginosa*. DM created and characterized the medium and low strength Anderson promoter – tsP constructs in the pSB1C3 plasmid and optimized the lysis assay. EM and MA created and characterized the first Anderson promoter – chromoprotein constructs under the high copy number origin of replication. MA produced the chromoprotein expression results found in Figure 1. LRL is a major contributor to the figure design, writing of the manuscript, and produced the initial protocol for the lysis assay. JDS is the primary contributor to the writing of the manuscript, analyzed the chromoprotein expression in cell and after lysis results, analyzed the Sanger sequencing results, and constructed all the figures and tables in the manuscript.

## Acknowledgements

This work was supported by NSF award 2031102. We acknowledge Renee Harper and the student workers in the William Jewell College department of Biology for preparation of basic reagents and supplies and the Liberty Wastewater Treatment Plant in Liberty, MO, for access to wastewater effluent for sampling purposes.

